# Altered Affective Behaviors in Casein Kinase 1 Epsilon Mutant Mice

**DOI:** 10.1101/2020.06.26.158600

**Authors:** Lili Zhou, Karrie Fitzpatrick, Christopher Olker, Martha H. Vitaterna, Fred W. Turek

## Abstract

Affective behaviors and mental health are profoundly affected by disturbances in circadian rhythms. Casein kinase 1 epsilon (CSNK1E) is an essential component of the core circadian clock. Mice with *tau* or null mutation of this gene have shortened and lengthened circadian period respectively. Here we examined anxiety-like, fear, and depressive-like behaviors in both male and female mice of these two different mutants. Compared with wild-type mice, we found reductions in fear and anxiety-like behaviors in both mutant lines and in both sexes, with the *tau* mutants exhibiting the greatest phenotypic changes. However, the depressive-like behaviors had distinct phenotypic patterns, with markedly less depressive-like behaviors in female null mutants, but not in *tau* mutants of either sex. To determine whether abnormal light entrainment of *tau* mutants to 24 hour light-dark cycles contributes to these phenotypic differences, we also examined these behaviors in *tau* mutants on a 20 hour light-dark cycle close to their endogenous circadian period. The normalized entrainment restored more wild-type-like behaviors for fear and anxiety, but it induced depressive-like behavior in *tau* mutant females. These data show that both mutations of *Csnk1e* broadly affect fear and anxiety-like behaviors, while the effects on depressive-like behavior vary with genetics, photoperiod, and sex, suggesting that the mechanisms by which *Csnk1e* affects fear and anxiety-like behaviors may be similar, but distinct from those affecting depressive-like behavior. Our study also provides experimental evidence in support of the hypothesis of beneficial outcomes from properly entrained circadian rhythms in terms of the anxiety-like and fear behaviors.

## Introduction

Circadian rhythms are endogenous oscillations present in diverse organisms with periodicities of approximately 24 hours. They can be synchronized with the external environment by both light and non-photic stimuli, such as feeding and social activities, allowing organisms to anticipate and prepare for the changes of environments, and thus may provide advantages for fitness and survival (Patke *et al*., 2019). A circadian resonance hypothesis has been proposed by Pittendrigh, one of the founding fathers of circadian biology, that the effects are beneficial to organisms when the external cycles are close to their evolved natural periods (Pittendrigh *et al*., 1959). Apart from ambiguous epidemiologic studies of prevalence of various symptoms and diseases in jet lag and shift workers, supportive evidence of this hypothesis is scattered and mainly focuses on reproductive fitness in cyanobacteria (Ouyang *et al*., 1998; Woelfle *et al*., 2004), flies (Pittendrigh & Minis, 1972; von Saint Paul & Aschoff, 1978; Beaver *et al*., 2002), and plants (Dodd *et al*., 2005). More recently, studies in rodents showed that the properly entrained rhythmicity led resistance to cardiorenal disease in hamsters (Martino *et al*., 2008), and had survival advantage in semi-natural conditions in mice (Spoelstra *et al*., 2016). However, no experimental evidence comes from mammalian models from the aspect of mental health.

Increasing evidence suggests that disturbances in circadian rhythms have profound consequences on affective states, cognition, and various other aspects of mental health (Boivin, 2000; Lamont *et al*., 2007; McClung, 2007; Turek, 2007). Many studies have found significant or suggestive associations between specific circadian genes and mood disorders, such as bipolar disorder, major depression, and seasonal affective disorder in the human population (Benedetti *et al*., 2003; Johansson *et al*., 2003; Mansour *et al*., 2006; Nievergelt *et al*., 2006; Partonen *et al*., 2007; Kripke *et al*., 2009; Mansour *et al*., 2009; McGrath *et al*., 2009; Severino *et al*., 2009; Lavebratt *et al*., 2010a; Lavebratt *et al*., 2010b; Soria *et al*., 2010; Terracciano *et al*., 2010; McCarthy *et al*., 2011; Liberman *et al*., 2017). Studies of mouse models have revealed altered affective behaviors as a result of disruptions in genes in the core circadian molecular pathway. For example, a mutation in *Per2* in mice causes both arrhythmic locomotor activity (Zheng *et al*., 2001) and a depression-resistant phenotype (Hampp *et al*., 2008). A mutation in the *Clock* gene (*Clock*^Δ*19*/Δ*19*^) that renders a lengthened circadian period and altered amplitude of rhythms (Vitaterna *et al*., 1994; Vitaterna *et al*., 2006) exhibit a wide range of behavioral abnormalities similar to that of mania including increased exploratory activity (Easton *et al*., 2003), decreased anxiety-like and depressive-like behaviors (Roybal *et al*., 2007), and dopaminergic hyperactivity in the ventral tegmental area (McClung *et al*., 2005). An ENU-induced mutation in *Fbxl3* gene resulted in lengthened circadian period (Godinho *et al*., 2007), and reduced anxiety- and depressive-like behaviors (Keers *et al*., 2012). Transgenic mice bearing rare human PER3 variants, P415A and H417R, which cause familial advanced sleep phase syndrome, or *Per3* null mutant mice were found to show depressive-like behaviors (Zhang *et al*., 2016). The knockdown *Bmal1* expression restricted to the SCN could increase depressive- and anxiety-like behaviors in mice (Landgraf *et al*., 2016). CRY in nucleus accumbens was found to regulate depressive-like behaviors in mice (Porcu *et al*., 2020).

Casein kinase 1 epsilon (CSNK1E) is an essential component of the core molecular circadian clock, which is wildly expressed in mouse nervous system (Magdaleno *et al*., 2006; Diez-Roux *et al*., 2011). The CSNK1E protein can phosphorylate a large number of substrates, including transcription factors, cytoskeletal proteins, and receptors, in the regulation of a wide range of physiological processes and diseases conditions (Knippschild *et al*., 2005). In circadian rhythms, CSNK1E phosphorylates PER and CRY proteins, the negative regulators in the central feedback loop of circadian clocks, thus facilitating their degradation and translocation, and consequently regulates the circadian period (Ko & Takahashi, 2006). In mice, the *tau* mutation of *Csnk1e* is a gain-of-function allele produced by insertion of a point mutation in the catalytic domain of CSNK1E (Meng *et al*., 2008). It accelerates the degradation of the PER proteins, causing homozygous mutants to exhibit a severely shortened circadian period of approximately 20 hours in the absence of the entraining effects of light (*i.e*., in constant darkness) (Ralph & Menaker, 1988; Lowrey *et al*., 2000; Meng *et al*., 2008). In contrast, when *Csnk1e* is knocked out, the null mutant mice have a slightly but significantly longer circadian period than wild-type mice (Meng *et al*., 2008).

Certain variants in *CSNK1E* have also been associated with an increased susceptibility to bipolar disorder (Shi *et al*., 2008). However, little is known about the effects of CSNK1E on psychiatric-related behaviors outside of this human association study, and no studies have been done in CSNK1E animal models. Given the critical role of CSNK1E in circadian rhythms, as well as the relationship between circadian rhythm disturbances and a variety of mood and cognition disorders, it is important to explore a broad spectrum of affective behavioral functions for CSNK1E. Therefore, we examined both *Csnk1e* null and *tau* mutant mice in a series of tests measuring anxiety-like, conditioned fear learning, and depressive-like behaviors. We included both males and females in this study to evaluate the sex effects on behaviors. Furthermore, housing *tau* mutants, which have severely shortened circadian period, under regular 24-hour light-dark (24-h LD) cycles essentially exposure them to long photoperiods. Thus, to investigate the extent to which abnormal entrainment to a 24-h LD cycle of *tau* mutants may affect affective behaviors, and to address the important circadian resonance hypothesis in terms of affective behaviors, a subset of *tau* mutants were tested in a 20-h LD cycle to permit normal light entrainment. Our study revealed significant effects of *Csnk1e* on affective behaviors in mice, and provides evidence for the circadian resonance hypothesis in anxiety-like and fear behaviors.

## Materials and Methods

### Mice

All mice used in the present study were housed and handled according to US Animal Welfare Act, and all procedures were approved in advance by the Institutional Animal Care and Use Committee at Northwestern University (Assurance Number: A3283-01).

The *Csnk1e^-/-^* (hereafter null) and *Csnk1e^tau/tau^* (hereafter *tau*) mutant mice used in the present study were created on a C57BL/6 background as described previously (Meng *et al*., 2008), and originated from homozygote breeding pairs obtained from the University of Manchester. Using these homozygote mice, new heterozygote mice were created at Northwestern University. And then, male and female null and *tau* mice and their wildtype littermate mice were all produced via the new heterozygote breeding pairs and maintained at Northwestern University. While undergoing the experiments described herein, wild-type and null mutant mice were maintained on a 12h:12h LD cycle (lights on at 0600h, lights off at 1800h), while the *tau* mutant mice were maintained on either a 12h:12h (24-h LD cycle) or a 10h:10h (20-h LD cycle) cycle. All *tau* mutant mice were originally bred and maintained on a 24-h LD cycle. At six to seven weeks of age (>10 days before behavioral tests), the LD cycle was changed to a 20-h LD cycle for the *tau* mice in the entrainment experiment (*tau* mutants were stably entrained to 10h:10h LD cycle within 2-3 days) until all behavioral tests were finished. All experiments were conducted at Zeitgeber Time (ZT0 = lights on) ZT4-ZT8. All mice were group housed with one to four same-sex littermates, with food and water available *ad libitum*. Light intensity and temperature remained constant at approximately 300 lux and 23 ± 2 °C, respectively. Male and female mice of two to four months of age were used for the behavioral tests, and the sample sizes for each of the behavioral tests ranged from N = 7 to 19 for males and N = 9 to 14 for females of each genotype.

### Elevated Plus Maze

The elevated plus maze (EPM) is a classic test utilized for assessing anxious behaviors, and consists of two open arms (each 57.8 cm long and 5.7 cm wide) alternating with two closed arms (each also 57.8 cm long and 5.7 cm wide, with 14.0 cm high walls on the three outer edges) situated in a plus formation, elevated by 30.5 cm. Mice were placed in the center of the maze and allowed to explore for 5 minutes, during which they were monitored from above by a video camera connected to a computer running video tracking software (Limelight v2.57, Actimetrics, Wilmette, IL). The mazes were cleaned with a 70% ethanol solution and allowed to dry between mice. The software tracks a midpoint over the mouse’s shoulders, which is used to determine the total distance traveled over the course of the test, as well as the number of entries into and the percentage of time spent in each arm of the maze, which were subsequently used to calculate the ratio of entries and time for the open versus closed arms. Reduced time spent in the open arms relative to wild types is indicative of an anxious phenotype.

### Open Field Activity

Assessment of open field activity (OFA) is also a commonly utilized test for the measurement of anxiety-like behavior. The apparatus used for measuring OFA consists of a 37.5 by 37.5 cm open arena surrounded by 36.4 cm tall walls. Mice were placed in the center of the open area and allowed to explore for 5 minutes, during which they were monitored from above by a video camera and tracking software as described above. The apparatus was cleaned with a 70% ethanol solution and allowed to dry between mice. The software package subdivides the arena into a grid of 5×5 sections of equal size, from which the nine inner-most sections are defined as the center, and the four sections abutting adjoining walls are defined as the corners. Tracking a midpoint over the mouse’s shoulders, the software determines the total distance traveled over the course of the test, as well as the number of entries into and the percentage of time spent in each of the defined subsections of the arena. Smaller amounts of time spent in or entries into the center area relative to wild types are indicative of an anxious phenotype.

### Fear Conditioning Test

The fear conditioning test (FCT) assesses learning and memory as well as fearful behaviors, which have been associated with multiple psychiatric disorders including anxiety disorders and schizophrenia. The FCT protocol used in the present study consisted of an 8 min test that occurred over two consecutive days. The test was conducted using individual closed, lighted chambers with metal grid floors (28.6 x 24.1 x 23.2 cm, Med Associates Inc., St. Albans, VT) that were each monitored by dedicated video cameras. On test Day 1, mice were first transported to a preparation room which was next to the test room, and allowed to habituate for 60 min in their home cages. Immediately before starting the test, mice were then brought into the dim FCT room, and were placed into the individual FC chambers, each containing an odor cue consisting of filter paper soaked with 5 mL of lemon oil. Mice were given 180 seconds to investigate the chamber with no applied stimuli. Immediately following this period, a 30-second auditory tone (2800 Hz) was presented with the last two seconds containing a simultaneous 0.75 mA shock delivered through the metal grid floor. This cue-shock combination was repeated three more times for a total of four presentations, each separated by 60-second periods of no applied stimuli. Chambers were cleaned with a 70% ethanol solution between mice. On test Day 2, the same procedure was conducted as in Day 1.

The fear response is quantified by the expression of freezing behavior, which is defined as the absence of movement other than respiration. Freezing is scored via a software package (FreezeFrame, Actimetrics, Wilmette, IL) calibrated to this motion threshold and reported in terms of the percentage of time spent freezing in the defined interval. Acquired fear was measured by the amount of freezing behavior during each 30-second period of auditory tone, with a lower level of freezing equated a reduction in fear acquisition. The conditioned contextual fear response was calculated by subtracting the amount of naïve baseline freezing behavior displayed during the first 180 seconds of initial exposure to the test chamber from that displayed during the first 180 seconds when the animal was returned to the test chamber one LD cycle later, such that a smaller difference equated a smaller conditioned contextual fear response.

To assess the sensitivity in response to electric shock, a separate group of mice with different genotypes were given a range of ascending electric stimulus (2mA-6.5mA). Behaviors in response to the stimulus were recorded by a video camera and tracking software for further analysis. The lowest electric current of the stimulus which evoked behaviors of jumping, vocalizing, and rearing was referred as the threshold of sensitivity to the stimulus.

### Tail Suspension Test

The tail suspension test (TST) was first designed as a method for screening antidepressants in mice, and it has been a standard and pharmacologically validated screen for measuring both antidepressant and depressive-like behavior. Mice were placed in a closed, lighted chamber suspended by the end of their tail with a custom modified clip so that the nose was 1-3 cm from the floor of the chamber, for six minutes. Sessions were recorded with a video camera mounted behind the chamber, and video files were scored via visual inspection at a one-second resolution. Immobility was defined as no voluntary body or limb movements, and was quantified in total seconds. Because differences in total immobility time can be accomplished by changes in both the number or the duration of the bouts if immobility behavior, the average duration as well as the total number of bouts of immobility were also calculated for each animal to examine the detailed struggling pattern. The chamber and clip were cleaned with a 70% ethanol solution between mice. Increased immobility relative to that of wild types is indicative of higher level of depressive-like behavior.

### Experimental order

To control carry-over effects, tests were consistently conducted in a specific order throughout different cohorts of mice as the following: EPM, OFA, FCT, and TST. Behaviors in the EPM and OFA were assessed on sequential days in the first week of testing, while the FCT was administered in the second week, and the TST was administered in the third week of testing. All tests were conducted during the light period between 5 and 10 hours after lights on in the 24-h LD condition, or between 5 and 8 hours after lights on in the 20-h LD condition. Experimenters were blind to the genotypes of the mice both during testing and the subsequent scoring of the recorded behaviors.

### Statistical Analysis

All statistical analyses and figures were produced using R (v 4.0.0) software (R Development Core Team, 2011). Comparisons between the genotypes and sexes were executed via two-way analysis of variance (ANOVA) tests, with Tukey HSD post-hoc tests performed when appropriate. The alpha level significance threshold was defined as p < 0.05. Principal component analysis (PCA) was performed using the “*prcomp*” function within the “*stats*” package available in R. Because different behavioral traits were measured in different units and on different scales, the data were standardized by mean-centering (mean = 0) and scaling to unit variance (variance = 1). In total, 13 behavioral measurements from the present study were subjected to PCA. The magnitude of the loadings (correlations) and their relative signs (+ or −) describe the influence of different traits on the principal components (PCs). To determine the optimal number of components to include in the model, the cross-validation method was performed using the “*Q2*” function within the “*pcaMethods*” package in R (Stacklies *et al*., 2007), where the peak value of the predicted variation, Q^2^, determines the optimal number of components to include in the model.

## Results

### *Csnk1e* mutations decrease anxiety-like behaviors in the EPM

Overall, *Csnk1e* mutations significantly altered anxiety-like behaviors as measured by the time spend in the open arms (*F*_3,81_ = 6.178, *P* = 0.0008) and numbers of entries in the open arms (*F*_3,81_ = 13.062, *P* < 0.0001) in the EPM (Figure. 1). In particular, under the 24-h LD condition, both null and *tau* mutant male and female mice spent a significantly higher percentage of time in the open arms of the maze than wild types (males: WT = 13.94 ± 1.48 %, null = 21.21 ± 2.45 %, *P* = 0.049, *tau* = 28.23 ± 3.34 %, *P* = 0.0005; females: WT = 15.22 ± 2.06 %, null = 22.30 ± 3.02 %, *P* = 0.048, *tau* = 37.31 ± 5.20 %, *P* = 0.003; Figure. 1A, B), with *tau* mutant mice showing a larger effect size than null mutant mice. A similar pattern was observed for the number of entries to open arms (males: WT = 14.41 ± 1.18, null = 19.64 ± 1.34, *P* = 0.04, *tau* = 24.67 ± 1.91, *P* = 0.0006; females: WT = 14.00 ± 1.89, null = 18.67 ± 1.79, *P* = 0.045, *tau* = 26.22 ± 1.96, *P* = 0.0002; Figure. 1C, D). Housing mice of 20-h circadian period under 24-h LD condition may abnormally entrain them. Therefore, in order to restore the environment more closely matching their endogenous circadian rhythms, the *tau* mutant mice were placed on a 20-h LD cycle. Interestingly, this resulted in a reduction of phenotype severity, restoring the altered behaviors toward wild type levels. Particularly, male *tau* mutant mice on a 20-h LD cycle were not significantly different from wild types in measures of the time spent in open arms (WT = 13.94 ± 1.48 %, *tau20h* = 18.30 ± 4.31 %, *P* = 0.54; Figure. 1A), and the numbers of entries to the open arms (WT = 14.41± 1.18, *tau*20h= 15.57 ± 1.52, *P* = 0.87; Figure. 1C). Although the female *tau* mutant mice on a 20-h LD cycle still showed significantly lower anxiety-like behaviors compared with wild types (time in open arms: WT = 15.22 ± 2.06 %, *tau20h* = 27.71 ± 3.24 %, *P* = 0.02; entries in open arms: WT = 14.00 ± 1.89, *tau*20h= 19.81 ± 1.52, *P* = 0.02; Figure. 1B, D), the effect size and significance level were lower than those of *tau* mutants on a 24-h LD cycle, suggesting a partial restoration of the phenotype. No significant differences were observed in the total distance traveled in the EPM among any of the groups and there were no significant differences between sexes for any of the EPM measures as indicated by ANOVA (data not shown).

**Figure 1.**
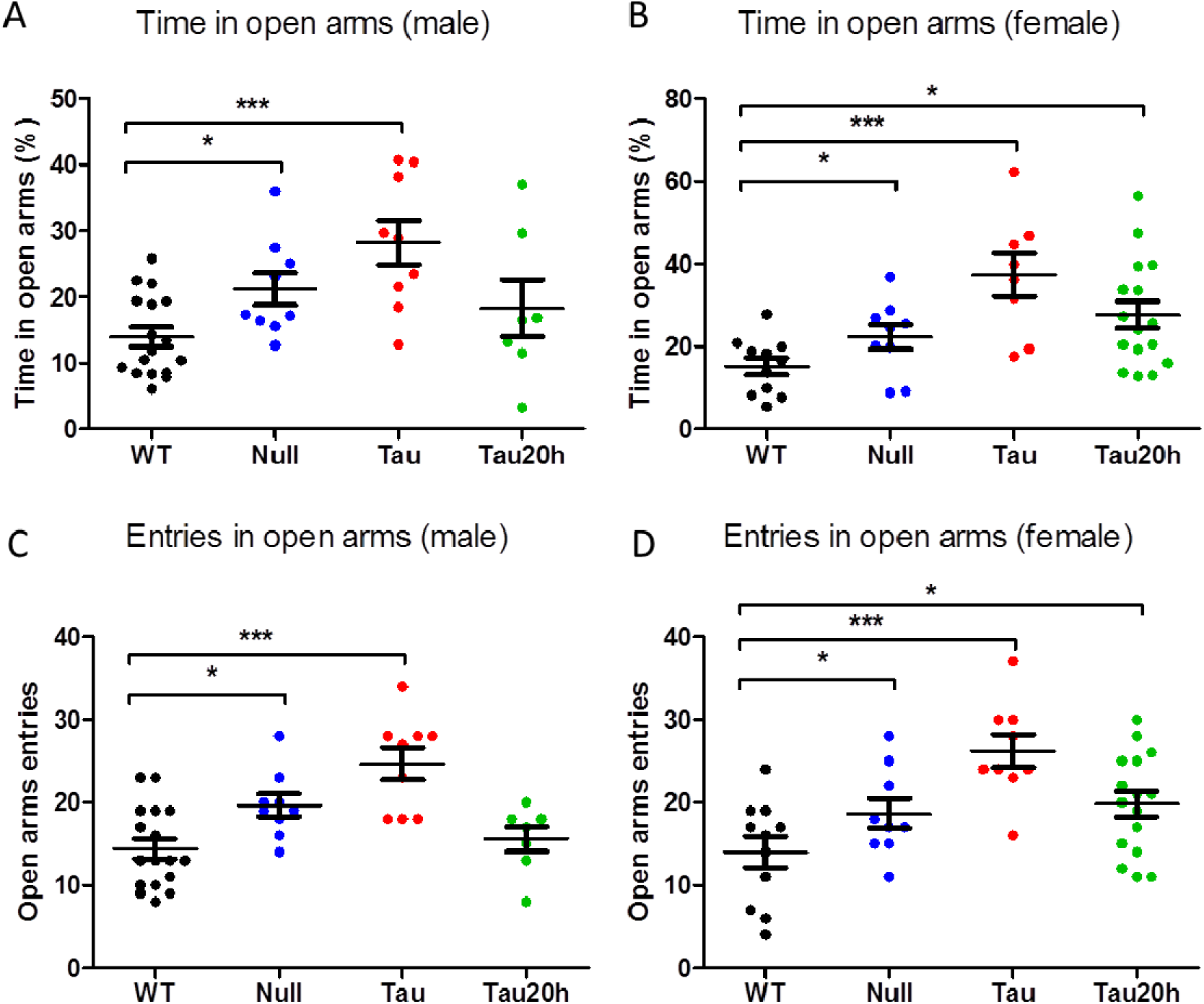
*Csnk1e* mutations decreased anxiety-like behaviors in the EPM. Percentage of total time spent in open arms in males (A) and females (B). Number of entries to open arms in males (C) and females (D). Data are presented as mean ± SEM. WT = *Csnk1e^+/+^*; Null = *Csnk1e^-/-^*; Tau = *Csnk1e^tau/tau^*; and Tau20h represents the *tau* genotype mice on a 20-h LD cycle, whereas the remaining groups were on a 24-h LD cycle. Asterisks denote significant differences between mutant genotype and wild type controls, \**P* < 0.05, \*\**P* < 0.01, \*\*\**P* < 0.001.

### *Csnk1e* mutations decreased anxiety-like behaviors in the OFA

Similar to EPM, the OFA is another commonly used test to measure anxiety-like behaviors in animal models. As expected, we observed similar behavioral patterns in the mutant mice in OFA as in EPM. Under 24-h LD cycle condition, both null and *tau* mutant mice exhibited a lower level of anxiety-like behaviors than wild type mice, as exhibited by a higher percentage of time spent in the center of the open field (males: WT = 11.03 ± 1.18 %, null = 14.76 ± 1.23 %, *P* = 0.038, *tau* = 16.04 ± 2.72 %, *P* = 0.049; females: WT = 10.10 ± 1.33 %, null = 16.56 ± 2.33 %, *P* = 0.032, *tau* = 16.64 ± 2.08 %, *P* = 0.019; Figure. 2A, B), and a higher number of entries into the center area of the open field (males: WT = 49.67 ± 4.82, null = 71.60 ± 7.66, *P* = 0.027, *tau* = 82.33 ± 8.35, *P* = 0.004; females: WT = 47.91 ± 6.38, null = 75.33 ± 9.83, *P* = 0.034, *tau* = 82.11 ± 8.85, *P* = 0.006; Figure. 2 C, D). However, when placed on a 20-h LD cycle, the behaviors of the *tau* mutant mice were restored to that of wild types, such that there were no more significant differences from wild types in any of the OFA measures in either sex (time in center: males: WT = 11.03 ± 1.18 %, *tau20h* = 8.69 ± 1.56 %, *P* = 0.25; females: WT = 10.10 ± 1.33 %, *tau*20h = 12.44 ± 1.23 %, *P* = 0.50; center entries: males: WT = 49.67 ± 4.85, *tau*20h= 41.71 ± 6.11, *P* = 0.32; females: WT = 47.91 ± 6.38, *tau*20h= 66.06 ± 6.31, *P* = 0.06; Figure. 2). The total distances mice traveled in the open field were not different between genotypes, and there were no significant differences between sexes for any of the OFA measures (data not shown).

**Figure 2.**
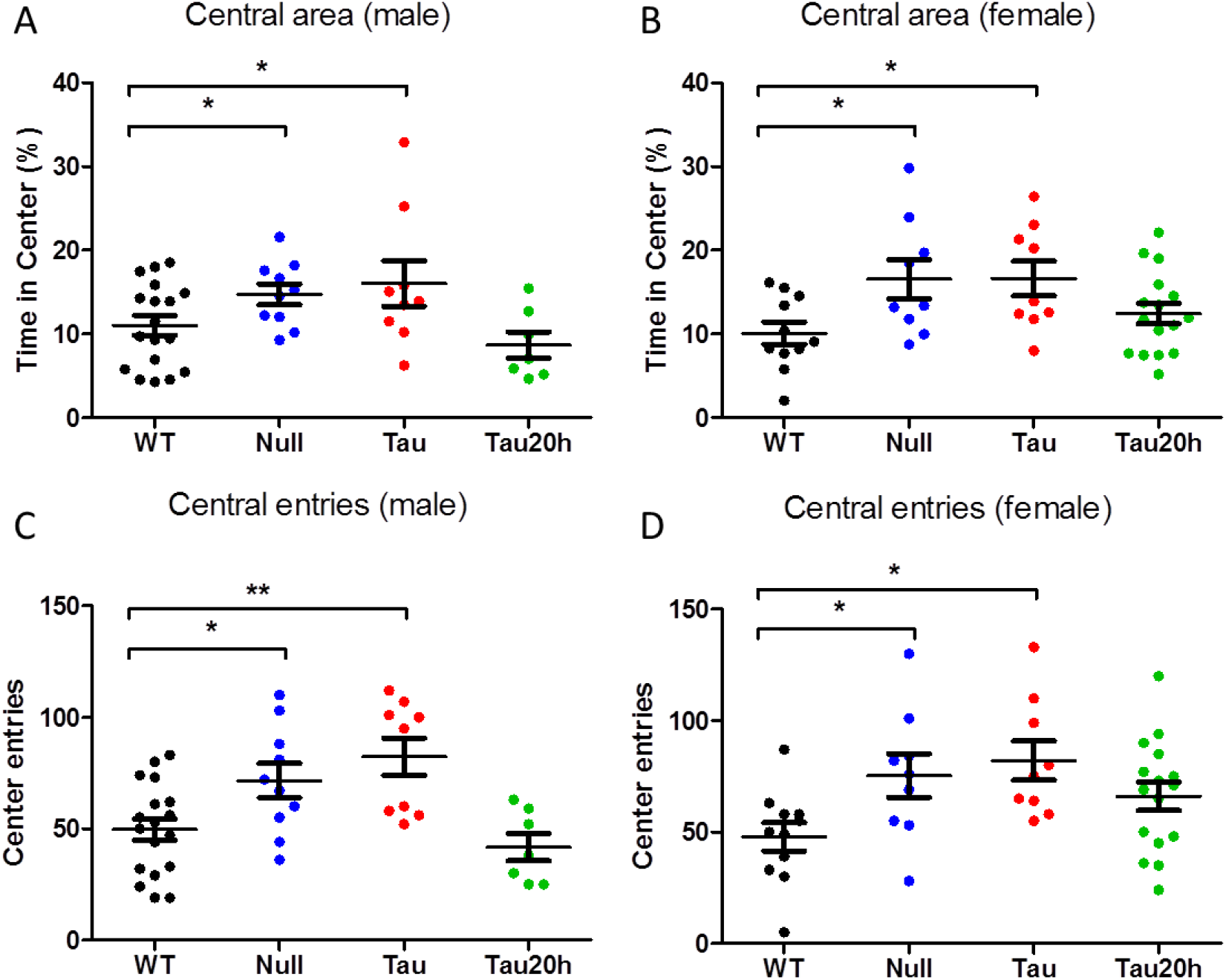
*Csnk1e* mutations decreased anxiety-like behaviors in the OFA. Percentage of time spent in the center area of the open field in males (A) and in females (B).Number of entries to the center area in males (E) and in females (F). Data are presented as mean ± SEM. WT = *Csnk1e^+/+^*; Null = *Csnk1e^-/-^*; Tau = *Csnk1e^tau/tau^*; and Tau20h represents the *tau* genotype mice on a 20-h LD cycle, whereas the remaining groups were on a 24-h LD cycle. Asterisks denote significant differences between mutant genotype and wild type controls, \**P* < 0.05, \*\**P* < 0.01, \*\*\**P* < 0.001.

### Impaired fear conditioning of *Csnk1e* mutants in the FCT

The *Csnk1e* mutant mice exhibited an impaired fear acquisition and contextual fear response compared with wild types in the FCT (Figure. 3). Under 24-h LD condition, both null and *tau* mutant males exhibited significantly lower fear acquisition than wild types during the conditioning training (Figure. 3A, B). A similar observation was found in *tau* mutant females under 24-h LD condition (Figure. 3E), though not in null mutant females (Figure 3D). However, when placed on a 20-h LD cycle, *tau* mutant mice exhibited normal fear acquisition as wild type mice in either sex (Figure. 3C, F). Twenty-four hour later on the day of fear test, both mutants and both sexes exhibited significant lower contextual fear responses than wild type mice under 24-h LD condition (males: WT = 29.42 ± 3.41 %, null = 17.78 ± 3.15 %, *P* = 0.043, *tau* = 11.24 ± 3.99 %, *P* = 0.01; females: WT = 30.98 ± 4.66 %, null = 16.38 ± 4.38 %, *P* = 0.03, *tau* = 8.64 ± 2.93%, *P* = 0.0009; Figure.3G, H). The 20-h LD cycle restored the contextual fear freezing to wildtype level in *tau* mutant males (WT = 29.42 ± 3.41%, *tau*20h = 21.46 ± 2.40 %, *P* = 0.29; Figure3. G), but not the in *tau* mutant females (WT = 30.98 ± 4.66 %, *tau*20h = 13.47 ± 2.30 %, *P* = 0.01; Figure3. H). The impaired fear acquisition and response in null and *tau* mutants was not due to their defective sensation to stimuli, because all mutant mice had a similar sensitivity to electric shock (*F*_2,81_ = 0.09, *P* = 0.91; Figure. 3I). The sex effect and interaction between sex and genotype were not significant in the FCT measures (data not shown).

**Figure 3.**
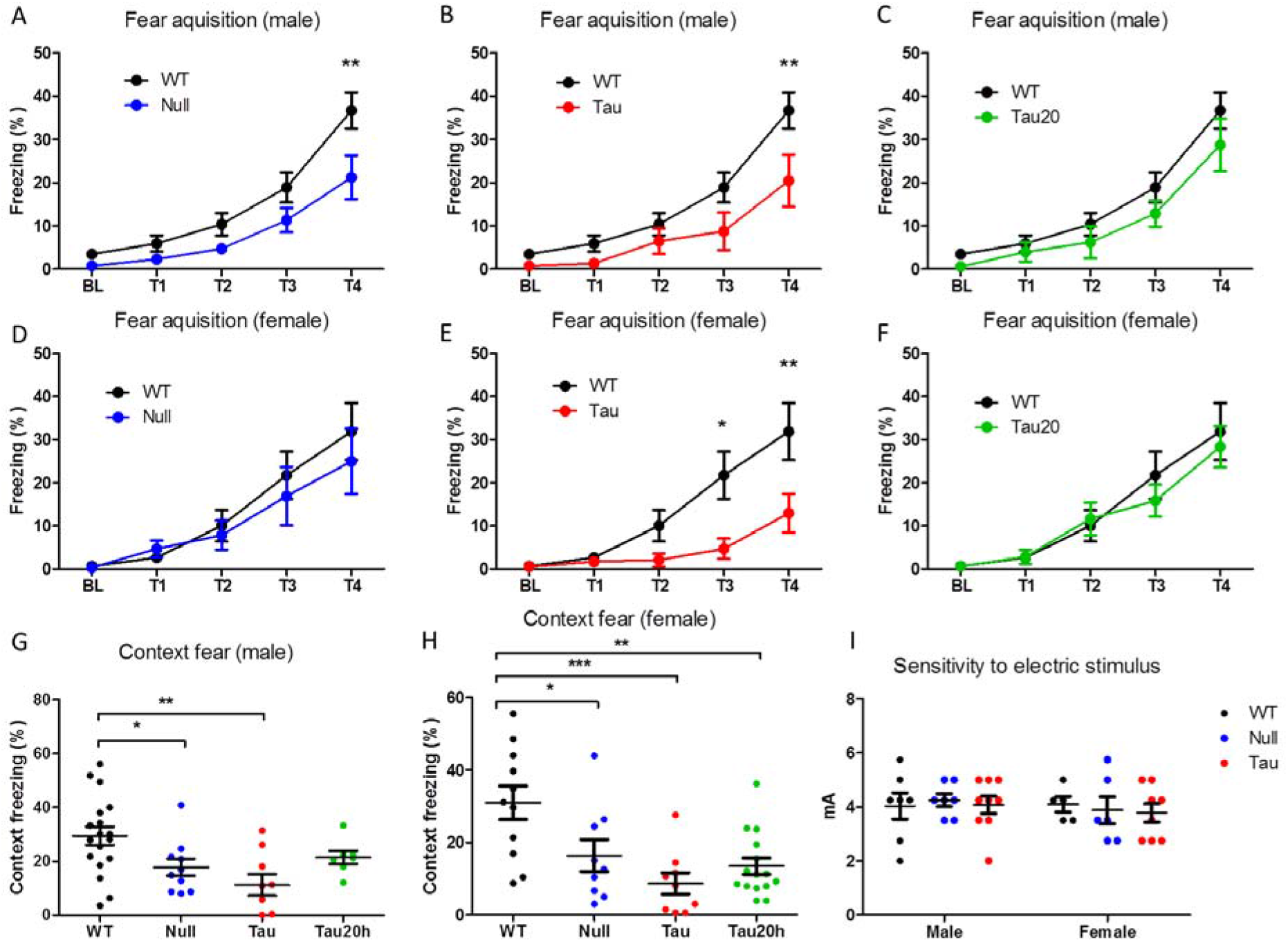
Fear conditioning was impaired in *Csnk1e* mutants in the FCT. Fear acquisition was measured as the percentage of freezing during the training of fear conditioning with four presentations of tone-shock pairings (T1-T4) in males (A-C) and females (D-F). BL represents the pre-training baseline level. Twenty-four hours later, the contextual fear response was measured as an increase in the percentage of freezing over baseline freezing in males (G) and females (H). (I) The sensitivity in response to electric stimuli were not significantly different between genotypes and sexes. Data are presented as mean ± SEM. WT = *Csnk1e^+/+^*; Null = *Csnk1e^-/-^*; Tau = *Csnk1e^tau/tau^*; and Tau20h represents the *tau* genotype mice on a 20-h LD cycle, whereas the remaining groups were on a 24-h LD cycle. Asterisks denote significant differences between mutant genotype and wild type controls, \**P* < 0.05, \*\**P* < 0.01, \*\*\**P* < 0.001.

### Depressive-like behaviors were altered by *Csnk1e* mutations in TST

The different *Csnk1e* mutations also altered depressive-like behaviors in the TST. We found significant genotype by sex interactions in measures of total immobility (*F*_3,81_ = 10.24, *P* < 0.001) and average immobility bout duration (*F*_3,81_ = 5.13, *P* = 0.003). *Csnk1e* null mutant mice exhibited generally lower levels of depressive-like behavior, with null mutant males displaying significantly lower average immobility bout durations than wild types (WT = 11.78 ± 1.23 s, null = 5.70 ± 1.05 s, *P* = 0.01; Figure. 4C), and null mutant females showing significantly less time staying immobility (WT = 54.82 ± 10.43 s, null = 32.00 ± 7.6 s, *P* = 0.02; Figure. 4B) and lower average immobility bout durations (WT = 8.69 ± 1.46 s, null = 4.26 ± 0.79 s, *P* = 0.02; Figure. 4D). For the *tau* mutant mice, however, the levels of the depressive-like behavior varied with both LD cycle and sex. Neither male nor female *tau* mutant mice were significantly different from wild types when placed on a 24-h LD cycle (Figure. 4). But when placed on a 20-h LD cycle, female *tau* mutant mice displayed significantly higher level of depressive-like behaviors, as measured by more time staying immobility (WT = 54.82 ± 10.43 s, *tau20h* = 92.63 ± 12.63 s, *P* = 0.048; Figure. 4B), and longer average immobility bout duration than those of wild types (WT = 8.69 ± 1.46 s, *tau*20h = 13.66 ± 1.12 s, *P* = 0.013; Figure. 4D). It is worth noting that the altered time in total immobility in mutants was due to the altered length of bout duration of immobility, rather than the number of bouts of immobility, as there were no significant differences between sexes, genotypes, and their interactions (Figure. 4E, F).

**Figure 4.**
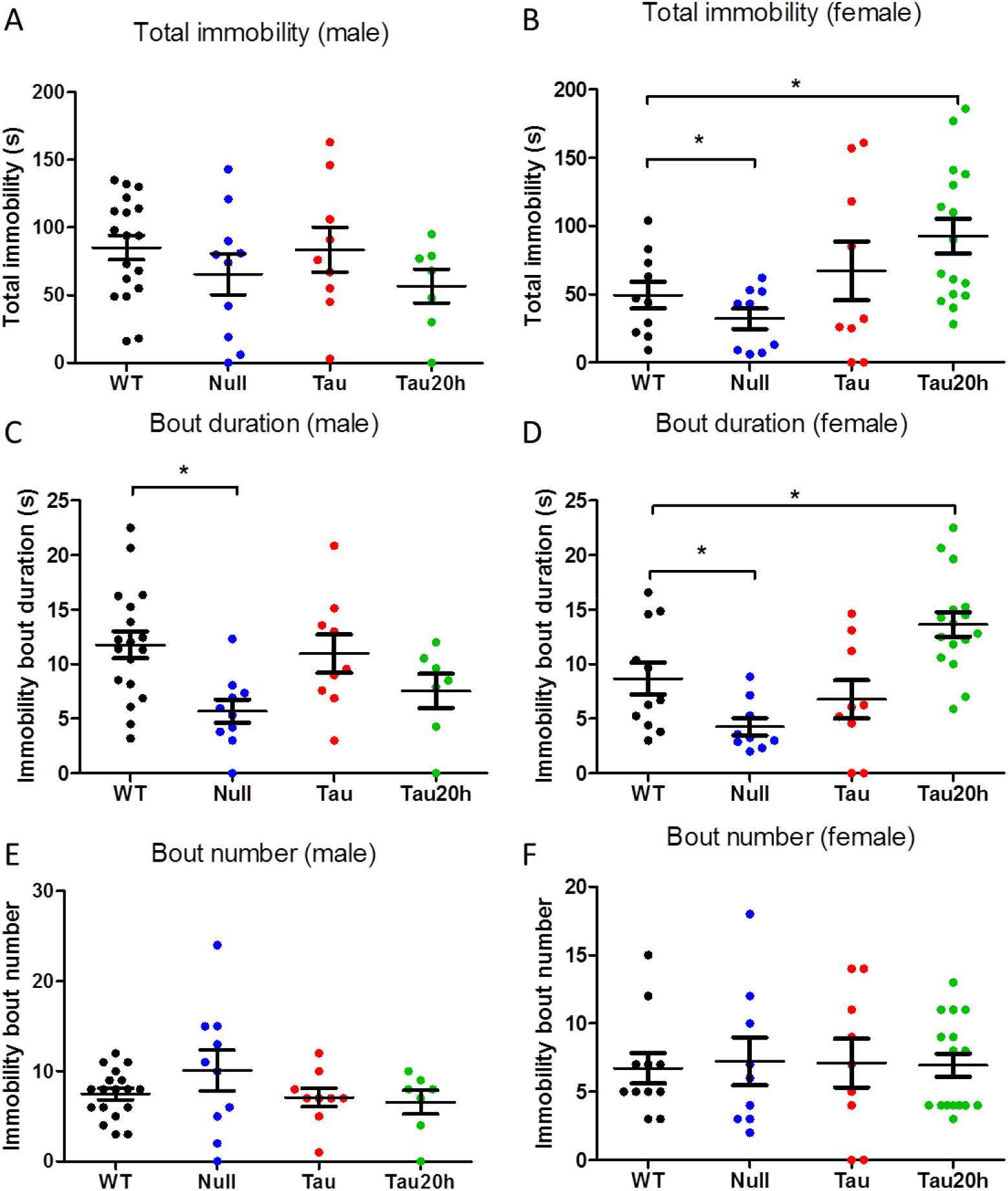
Depressive-like behaviors were altered by *Csnk1e* mutations in TST. Total immobility (seconds) in males (A) and females (B). Average immobility bout duration in males(C) and females (D). Number of immobility bout in males (E) and females (F). Data are presented as mean ± SEM. WT = *Csnk1e^+/+^*; Null = *Csnk1e^-/-^*; Tau = *Csnk1e^tau/tau^*; and Tau20h represents the *tau* genotype mice on a 20-h LD cycle, whereas the remaining groups were on a 24-h LD cycle. Asterisks denote significant differences between mutant genotype and wild type controls, \**P* < 0.05, \*\**P* < 0.01, \*\*\**P* < 0.001.

### PCA revealed depressive-like behavior as a distinct cluster

To test the hypothesis that depressive-like behavior is separable from fear and anxiety-like behaviors in *Csnk1e* mutant mice, PCA was performed using 13 behavioral measurements from the four different behavioral tests performed in the study. The first three principal components were selected because they explained the maximum predicted variation (Figure. 5A), 64.2% of the variation in sum. PCA revealed that the depressive-like behavior was separate from the other behaviors, and primarily accounts for the second PC as determined by the large loading values of TST measures into the second component (Figure. 5B-D).

**Figure 5.**
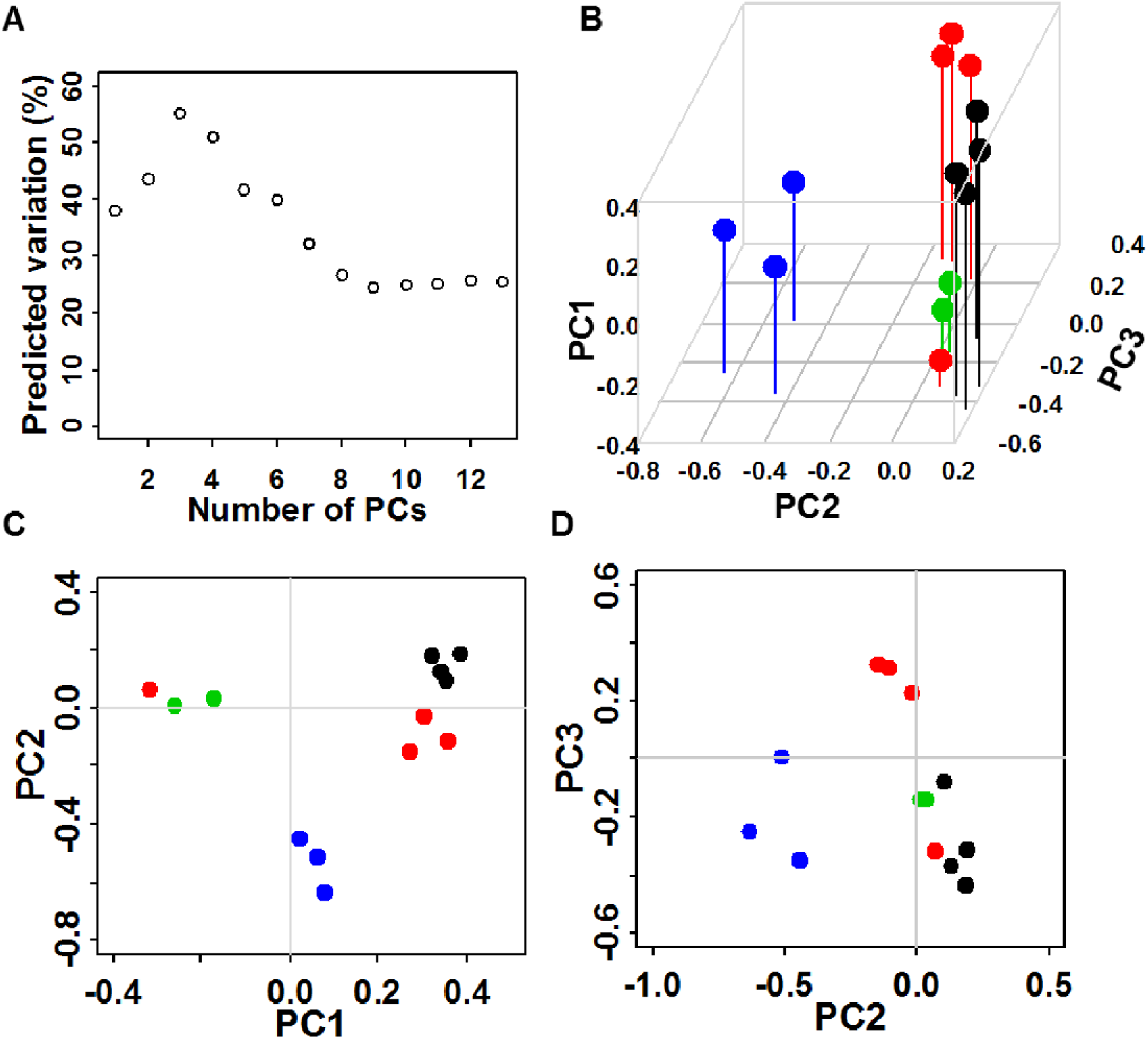
PCA revealed depressive-like behavior as a distinct cluster. (A) Plot of predicted variation (Q^2^). The optimal number of PCs was determined by estimating the predicted variation Q^2^ of the model with increasing number of components, where the peak of predicted variation indicates the optimal number of components that should go into the PC model. (B) Loading plot of the behavioral phenotypes in the first three principal components, explaining 64.2% of the variance. (C) Loading plot of the behavioral phenotypes in the first and the second principal components. (D) Loading plot of the behavioral phenotypes in the second and the third principal components. Colors indicate the behavioral tests: black = EPM, red = OFA, green = FCT, blue= TST.

## Discussion

The present study characterized anxiety-like, fear, and depressive-like behaviors of *Csnk1e* mutant mice through measuring their activity in series behavioral tests. We used two distinct genetically engineered mice carrying different mutations within the *Csnk1e* gene throughout the body: a gain-of-function *tau* mutation and a loss-of-function null mutation. *Csnk1e tau* mutant mice exhibit a significant 4-hr shortening of circadian period and abnormal phase angle of entrainment. In contrast, *Csnk1e* null mutant mice exhibit a very small but significant circadian period lengthening (Meng *et al*., 2008). We found that genetic alterations of *Csnk1e* have a robust effect on anxiety-like, fear, and depressive-like behaviors that varied in severity with the level of *Csnk1e* functionality. Additionally, we found that the disorganization of circadian rhythms caused by a misalignment between the LD cycle and endogenous circadian period has a strong impact on some of those behaviors.

Under 24-h LD entrained condition, both *tau* and null mutants exhibited lower levels of anxiety-like and fear behaviors than wild types, with the *tau* mutants showing larger effect size than the null mutants. Considering the opposing effects of these two mutations in circadian rhythms, it suggests that the function of CSNK1E on anxiety-like and fear behaviors to some extent might not be through the core circadian clock, such as CRY and PER proteins. The milder effect size in null mutants may be due to the compensation by the other kinase homologs, given that casein kinases are a large gene family. But this does not mean that the effects of CSKN1E on anxiety-like and fear behaviors are completely independent of circadian rhythms. As shown in this study, when the *tau* mutant mice were on a 20-h LD cycle, which corresponds with their endogenous circadian period and allows for normal entrainment, anxiety-like and fear behaviors were either restored to levels of wild types, as observed with OFA, or shifted in that direction, as seen in the EPM and FCT. Thus, when the *tau* mutant mice were able to entrain to LD cycle with a normal phase relationship, anxiety-like and fear behavioral alterations were either fully or partially rescued. Notably, in a study complementary to our work, wildtype mice housed under 20-h LD cycles showed decreased latency to enter a novel environment, suggesting lower fear/anxiety responses (Karatsoreos *et al*., 2011). Similar results have been reported with respect to cardiovascular and renal diseases in hamsters carrying the *tau* mutation. The incidence of cardiomyopathy and renal disease observed on 24-h LD cycles were absent in *tau* hamsters maintained on a LD cycle that was matched to their endogenous circadian period (Martino *et al*., 2008). These results support Pittendrigh’s hypothesis that maintaining synchrony between internal clocks and external zeitgebers is important for maintaining normal behavior and physiology.

Addressing Pittendrigh’s hypothesis is not trivial. It points to a central question of how important of our circadian rhythms from an evolutionary perspective. Unfortunately, there is still very little experimental evidence in support of this long standing hypothesis, especially in mammals. Despite the fantastic work showing natural selection against *tau* mutant allele in mice under semi-natural conditions (Spoelstra *et al*., 2016), it is still inconclusive about whether this was caused by circadian rhythm dysregulation or pleiotropic effect of *tau* allele. Converse experiments, in which *tau* and wild-type mice are exposed to short periods, are still needed. It might be noted that the circadian misalignment in our study resulted in lowered anxiety-like and fear behaviors. Superficially, this observation may be interpreted as disrupted circadian rhythms led to an anxiolytic “good outcome”, which seems contradictory to the notion that the effects are beneficial to organisms when properly entrained. However, from an evolutionary point of view, anxiety is an emotion associated with avoidance of danger and appropriate level of anxiety increases fitness in response to stress which may cause a loss of reproductive resources (Nesse, 1994). Too little anxiety/fear could result in higher risk to survival, which might be one possible reason for natural selection against the *tau* mutant allele in the previous study (Spoelstra *et al*., 2016). Thus, the lowered anxiety-like behavior in this study may be better interpreted as a deviation from the normal level.

In contrast, the *tau* and null mutations had distinct effects on depressive-like behavior. Only null mutants showed antidepressant-like phenotype, whereas *tau* mutants had similar level depressive-like behaviors to wild types. This suggests that the effector of *Csnk1e* on depressive-like behaviors may reside outside of the *tau* site, and be less likely compensated by other kinase homologs. Normalizing light entrainment was expected to benefit the behavioral outcome instead of increasing the severity, as seen in the anxiety-like and fear behaviors in this study. However, when the *tau* mutant mice were maintained on a 20-h LD cycle, the LD cycle that more closely matched their endogenous circadian period, actually induced depressive-like behaviors in female *tau* mutants. This unexpected result suggests that depressive-like behavior might be more susceptible to the duration of light exposure instead of the light entrainment. In support of this hypothesis, the depressive-like behavior in *Per3* mutant mice was found to be exacerbated by subjecting animals to a short photoperiod (Zhang *et al*., 2016).

We also observed a significant interaction between sex and genotype, but only in depressive-like behavior. The male null mutants exhibited a more robust antidepressant-like phenotype than the female null mutants, and the female *tau* mutant mice displayed a depressive-like phenotype on the 20-h LD cycle while the male *tau* mutants did not. This suggests that the regulation of depressive-like behavior by *Csnk1e* might be prone to the influence of genetic sex or sex hormones. The sex-linked effect in depression is also seen in humans, as it is well-known that the incidence of depression varies by sex, with women having approximately twice the rate of major depression and 1.5 times the rate of mood disorders in general (Parker & Brotchie, 2010). Animal models of depression, as well as stress reactivity and other biomarkers associated with depressive-like behaviors, have also been demonstrated as having differing effects based on sex (Dalla *et al*., 2010; Dalla *et al*., 2011). Female animal models are underutilized in neuroscience research, and inclusion of female mice in research is important and desirable (Prendergast *et al*., 2014). Our results implicate both circadian rhythm dysregulation and *Csnk1e* in the sex-specific vulnerability to depressive-like behaviors.

In our study, two different mutations of *Csnk1e* resulted in similar phenotypic patterns in anxiety-like and fear behaviors, with the *tau* mutation resulted in more extreme phenotypes than the null mutation. However, these mutations resulted in different and more complex patterns in depressive-like behavior. PCA further indicated that depressive-like behavior was separate from the other behaviors examined in *Csnk1e* mutant mice in the present study. This gives rise to an interesting question about the underlying neurobiological components involved in regulating anxiety, fear, and depression behaviors. Fear and anxiety are generally considered separate behavioral conditions or reactions, with fear typically considered as a reaction to an explicit threat and anxiety as a more general and prolonged state of distress with less distinct cues (Lang *et al*., 2000). These two basic behaviors have long been studied and been observed to have multiple interactions, significant comorbidity, and overlapping symptomatology (Schwartz & Weinberger, 1980; Dobson, 1985; Gorman, 1996; Cerdá *et al*., 2010; Sokoloff *et al*., 2011). Supporting evidence includes overlapping neuroanatomical areas and features found to be involved in both fear and anxiety, as well as physiological responses and neurochemical associations, as seen in humans and rodents alike (Lang *et al*., 2000; LeDoux, 2000; Richardson *et al*., 2004). Furthermore, studies in both rats (Fernandez-Teruel *et al*., 2002) and mice (Ponder *et al*., 2007a; Ponder *et al*., 2007b; Sokoloff *et al*., 2011) have indicated that anxiety and learned fear share a genetic basis. On the other hand, while comorbidity rates are highest for major depressive disorder and anxiety disorders (Kessler *et al*., 2005), they are still regarded as separate disorder categories with multiple distinct behavioral and affective symptomatology (American Psychiatric Association, 2000). In the present study, we found that anxiety-like and fear behaviors elicited through contextual fear conditioning exhibited a similar pattern throughout multiple perturbations to the circadian system, in contrast to that of depressive-like behavior, suggesting that the underlying neurobiological mechanisms involved in the regulation of these interrelated behaviors may have different mechanisms of interaction with the circadian system for anxiety and fear than for those of depression behaviors.

CSNK1E is not the only kinase component in the circadian system that is involved in the regulation of affective behaviors. It has been shown that the mood stabilizer lithium can inhibit affective behaviors by way of inhibiting the action of the Glycogen synthase kinase 3 beta (GSK3B) (Quiroz *et al*., 2004; Gould & Manji, 2005). Transgenic mice overexpressing *Gsk3b* exhibited reduced levels of despair behaviors, as well as hyperactivity and an increased startle response (Prickaerts *et al*., 2006). Intriguingly, the activation of GSK3B can be positively modulated by CSNK1E (Svenningsson *et al*., 2003), suggesting possible shared pathways involving these two kinases for mood regulation. The closest isoform of *Csnk1e* is the Casein kinase 1 delta (*Csnk1d*), which is considered to have some redundancy with *Csnk1e* in many physiological processes, including circadian rhythms. Mice overexpressing *Csnk1d* exhibited increased locomotor activity and decreased anxiety-like behaviors in an open field paradigm (Zhou *et al*., 2010), which parallels the pattern reported here in *Csnk1e* mutant mice. Collectively, these findings lend further support to the involvement of *Csnk1e* mechanisms in the regulation of affective behaviors. It is noteworthy that kinases represent one of the largest groups of druggable targets in the human genome (Oprea *et al*., 2018). Although the links between circadian rhythms and affective behaviors have been revealed in many other circadian genes, the focus was primarily on transcription factors such as *Bmal1* and *Clock*. Given the potential therapeutic importance, kinases such as *Csnk1e* are worth gaining more attention.

In conclusion, the present study highlighted the deleterious effects of misaligned circadian rhythms and beneficial effects of properly synchronized circadian rhythms on anxiety-like and fear behaviors. It provides experimental evidence for Pittendrigh’s longstanding hypnosis. Our results demonstrated that the circadian clock gene *Csnk1e* was involved in the regulation of anxiety-like, fear, and depressive-like behaviors, which was influenced by circadian desynchrony and, in the case of depressive-like behaviors, sex. Results suggested that fear and anxiety were more closely linked behaviors and were more likely to have shared underlying molecular mechanisms. On the other hand, the regulation of depression might be more associated with circadian clock function as well as other factors such as genetic sex or sex hormones. These different animal models of *Csnk1e* provide valuable opportunities for dissecting the mechanisms underlying the circadian rhythms and different categories of affective disorders in the future. An improved understanding and pharmacological control of *Csnk1e* could have implications for the drug discovery and treatment of affective disorders in humans.

## Acknowledgments

We thank Dr. Kazuhiro Shimomura for his contributions in multiple discussions and helpful suggestions, and Dr. Andrew Loudon, Dr. Joseph Takahashi and Dr. Caroline Ko for kindly providing the *Csnk1e* mutant mice.

## Competing Interests

The authors declare no conflicts of interests.

## Author Contributions

LZ, MHV, and FWT designed the study; LZ, KF and CO conducted the experiments and collected the data; LZ analyzed the data; LZ wrote the manuscript; all authors revised and approved the manuscript.

## Data Accessibility

Data can be supplied from the corresponding authors on reasonable request.

## Abbreviations

ANOVA: Analysis of Variance
CSNK1D: Casein kinase 1 delta
CSNK1E: Casein kinase 1 epsilon
EPM: Elevated Plus Maze
FCT: Fear Conditioning Test
GSK3B: Glycogen synthase kinase 3 beta
LD: Light-Dark
OFA: Open Field Activity
PCA: Principal Component Analysis
TST: Tail Suspension Test
ZT: Zeitgeber Time

## Notes

### Competing Interest Statement

The authors have declared no competing interest.

## References

Association, A.P. (2000) Diagnostic and Statistical Manual of Mental Disorders, Fourth Edition: DSM-IV-TR^®^. American Psychiatric Association.

Beaver, L., Gvakharia, B., Vollintine, T., Hege, D., Stanewsky, R. & Giebultowicz, J. (2002) Loss of circadian clock function decreases reproductive fitness in males of Drosophila melanogaster. Proceedings of the National Academy of Sciences, 99, 2134–2139.

Benedetti, F., Serretti, A., Colombo, C., Barbini, B., Lorenzi, C., Campori, E. & Smeraldi, E. (2003) Influence of CLOCK gene polymorphism on circadian mood fluctuation and illness recurrence in bipolar depression. Am J Med Genet B Neuropsychiatr Genet, 123B, 23–26.

Boivin, D.B. (2000) Influence of sleep-wake and circadian rhythm disturbances in psychiatric disorders. J Psychiatry Neurosci, 25, 446–458.

Cerdá, M., Sagdeo, A., Johnson, J. & Galea, S. (2010) Genetic and environmental influences on psychiatric comorbidity: A systematic review. Journal of Affective Disorders, 126, 14–38.

Dalla, C., Pitychoutis, P.M., Kokras, N. & Papadopoulou-Daifoti, Z. (2010) Sex Differences in Animal Models of Depression and Antidepressant Response. Basic & Clinical Pharmacology & Toxicology, 106, 226–233.

Dalla, C., Pitychoutis, P.M., Kokras, N. & Papadopoulou-Daifoti, Z. (2011) Sex differences in response to stress and expression of depressive-like behaviours in the rat. Curr Top Behav Neurosci, 8, 97–118.

Diez-Roux, G., Banfi, S., Sultan, M., Geffers, L., Anand, S., Rozado, D., Magen, A., Canidio, E., Pagani, M., Peluso, I., Lin-Marq, N., Koch, M., Bilio, M., Cantiello, I., Verde, R., De Masi, C., Bianchi, S.A., Cicchini, J., Perroud, E., Mehmeti, S., Dagand, E., Schrinner, S., Nürnberger, A., Schmidt, K., Metz, K., Zwingmann, C., Brieske, N., Springer, C., Hernandez, A.M., Herzog, S., Grabbe, F., Sieverding, C., Fischer, B., Schrader, K., Brockmeyer, M., Dettmer, S., Helbig, C., Alunni, V., Battaini, M.A., Mura, C., Henrichsen, C.N., Garcia-Lopez, R., Echevarria, D., Puelles, E., Garcia-Calero, E., Kruse, S., Uhr, M., Kauck, C., Feng, G., Milyaev, N., Ong, C.K., Kumar, L., Lam, M., Semple, C.A., Gyenesei, A., Mundlos, S., Radelof, U., Lehrach, H., Sarmientos, P., Reymond, A., Davidson, D.R., Dollé, P., Antonarakis, S.E., Yaspo, M.L., Martinez, S., Baldock, R.A., Eichele, G. & Ballabio, A. (2011) A high-resolution anatomical atlas of the transcriptome in the mouse embryo. PLoS biology, 9, e1000582.

Dobson, K.S. (1985) The relationship between anxiety and depression. Clinical Psychology Review, 5, 307–324.

Dodd, A.N., Salathia, N., Hall, A., Kévei, E., Tóth, R., Nagy, F., Hibberd, J.M., Millar, A.J. & Webb, A.A. (2005) Plant circadian clocks increase photosynthesis, growth, survival, and competitive advantage. Science (New York, N.Y.), 309, 630–633.

Easton, A., Arbuzova, J. & Turek, F.W. (2003) The circadian Clock mutation increases exploratory activity and escape-seeking behavior. Genes Brain Behav, 2, 11–19.

Fernandez-Teruel, A., Escorihuela, R.M., Gray, J.A., Aguilar, R., Gil, L., Gimenez-Llort, L., Tobena, A., Bhomra, A., Nicod, A., Mott, R., Driscoll, P., Dawson, G.R. & Flint, J. (2002) A quantitative trait locus influencing anxiety in the laboratory rat. Genome Res, 12, 618–626.

Godinho, S.I., Maywood, E.S., Shaw, L., Tucci, V., Barnard, A.R., Busino, L., Pagano, M., Kendall, R., Quwailid, M.M., Romero, M.R., O’Neill, J., Chesham, J.E., Brooker, D., Lalanne, Z., Hastings, M.H. & Nolan, P.M. (2007) The after-hours mutant reveals a role for Fbxl3 in determining mammalian circadian period. Science (New York, N.Y.), 316, 897–900.

Gorman, J.M. (1996) Comorbid depression and anxiety spectrum disorders. Depress Anxiety, 4, 160–168.

Gould, T.D. & Manji, H.K. (2005) Glycogen synthase kinase-3: a putative molecular target for lithium mimetic drugs. Neuropsychopharmacology, 30, 1223–1237.

Hampp, G., Ripperger, J.A., Houben, T., Schmutz, I., Blex, C., Perreau-Lenz, S., Brunk, I., Spanagel, R., Ahnert-Hilger, G., Meijer, J.H. & Albrecht, U. (2008) Regulation of monoamine oxidase A by circadian-clock components implies clock influence on mood. Curr Biol, 18, 678–683.

Johansson, C., Willeit, M., Smedh, C., Ekholm, J., Paunio, T., Kieseppa, T., Lichtermann, D., Praschak-Rieder, N., Neumeister, A., Nilsson, L.G., Kasper, S., Peltonen, L., Adolfsson, R., Schalling, M. & Partonen, T. (2003) Circadian clock-related polymorphisms in seasonal affective disorder and their relevance to diurnal preference. Neuropsychopharmacology, 28, 734–739.

Karatsoreos, I.N., Bhagat, S., Bloss, E.B., Morrison, J.H. & McEwen, B.S. (2011) Disruption of circadian clocks has ramifications for metabolism, brain, and behavior. Proceedings of the national Academy of Sciences, 108, 1657–1662.

Keers, R., Pedroso, I., Breen, G., Aitchison, K.J., Nolan, P.M., Cichon, S., Nöthen, M.M., Rietschel, M., Schalkwyk, L.C. & Fernandes, C. (2012) Reduced anxiety and depression-like behaviours in the circadian period mutant mouse afterhours. PLoS One, 7, e38263–e38263.

Kessler, R.C., Chiu, W.T., Demler, O., Merikangas, K.R. & Walters, E.E. (2005) Prevalence, severity, and comorbidity of 12-month DSM-IV disorders in the National Comorbidity Survey Replication. Arch Gen Psychiatry, 62, 617–627.

Knippschild, U., Gocht, A., Wolff, S., Huber, N., Lohler, J. & Stoter, M. (2005) The casein kinase 1 family: participation in multiple cellular processes in eukaryotes. Cellular signalling, 17, 675–689.

Ko, C.H. & Takahashi, J.S. (2006) Molecular components of the mammalian circadian clock. Human Molecular Genetics, 15, R271–R277.

Kripke, D.F., Nievergelt, C.M., Joo, E., Shekhtman, T. & Kelsoe, J.R. (2009) Circadian polymorphisms associated with affective disorders. J Circadian Rhythms, 7, 2.

Lamont, E.W., James, F.O., Boivin, D.B. & Cermakian, N. (2007) From circadian clock gene expression to pathologies. Sleep Med, 8, 547–556.

Landgraf, D., Long, J.E., Proulx, C.D., Barandas, R., Malinow, R. & Welsh, D.K. (2016) Genetic Disruption of Circadian Rhythms in the Suprachiasmatic Nucleus Causes Helplessness, Behavioral Despair, and Anxiety-like Behavior in Mice. Biological psychiatry, 80, 827–835.

Lang, P.J., Davis, M. & Ã–hman, A. (2000) Fear and anxiety: animal models and human cognitive psychophysiology. Journal of Affective Disorders, 61, 137–159.

Lavebratt, C., Sjoholm, L.K., Partonen, T., Schalling, M. & Forsell, Y. (2010a) PER2 variantion is associated with depression vulnerability. Am J Med Genet B Neuropsychiatr Genet, 153B, 570–581.

Lavebratt, C., Sjoholm, L.K., Soronen, P., Paunio, T., Vawter, M.P., Bunney, W.E., Adolfsson, R., Forsell, Y., Wu, J.C., Kelsoe, J.R., Partonen, T. & Schalling, M. (2010b) CRY2 is associated with depression. PLoS One, 5, e9407.

LeDoux, J.E. (2000) Emotion circuits in the brain. Annu Rev Neurosci, 23, 155–184.

Liberman, A.R., Kwon, S.B., Vu, H.T., Filipowicz, A., Ay, A. & Ingram, K.K. (2017) Circadian Clock Model Supports Molecular Link Between PER3 and Human Anxiety. Scientific Reports, 7, 9893.

Lowrey, P.L., Shimomura, K., Antoch, M.P., Yamazaki, S., Zemenides, P.D., Ralph, M.R., Menaker, M. & Takahashi, J.S. (2000) Positional syntenic cloning and functional characterization of the mammalian circadian mutation tau. Science (New York, N.Y.), 288, 483–492.

Magdaleno, S., Jensen, P., Brumwell, C.L., Seal, A., Lehman, K., Asbury, A., Cheung, T., Cornelius, T., Batten, D.M., Eden, C., Norland, S.M., Rice, D.S., Dosooye, N., Shakya, S., Mehta, P. & Curran, T. (2006) BGEM: an in situ hybridization database of gene expression in the embryonic and adult mouse nervous system. PLoS biology, 4, e86.

Mansour, H.A., Talkowski, M.E., Wood, J., Chowdari, K.V., McClain, L., Prasad, K., Montrose, D., Fagiolini, A., Friedman, E.S., Allen, M.H., Bowden, C.L., Calabrese, J., El-Mallakh, R.S., Escamilla, M., Faraone, S.V., Fossey, M.D., Gyulai, L., Loftis, J.M., Hauser, P., Ketter, T.A., Marangell, L.B., Miklowitz, D.J., Nierenberg, A.A., Patel, J., Sachs, G.S., Sklar, P., Smoller, J.W., Laird, N., Keshavan, M., Thase, M.E., Axelson, D., Birmaher, B., Lewis, D., Monk, T., Frank, E., Kupfer, D.J., Devlin, B. & Nimgaonkar, V.L. (2009) Association study of 21 circadian genes with bipolar I disorder, schizoaffective disorder, and schizophrenia. Bipolar Disord, 11, 701–710.

Mansour, H.A., Wood, J., Logue, T., Chowdari, K.V., Dayal, M., Kupfer, D.J., Monk, T.H., Devlin, B. & Nimgaonkar, V.L. (2006) Association study of eight circadian genes with bipolar I disorder, schizoaffective disorder and schizophrenia. Genes Brain Behav, 5, 150–157.

Martino, T.A., Oudit, G.Y., Herzenberg, A.M., Tata, N., Koletar, M.M., Kabir, G.M., Belsham, D.D., Backx, P.H., Ralph, M.R. & Sole, M.J. (2008) Circadian rhythm disorganization produces profound cardiovascular and renal disease in hamsters. Am J Physiol Regul Integr Comp Physiol, 294, R1675–1683.

McCarthy, M.J., Nievergelt, C.M., Shekhtman, T., Kripke, D.F., Welsh, D.K. & Kelsoe, J.R. (2011) Functional genetic variation in the Rev-Erbalpha pathway and lithium response in the treatment of bipolar disorder. Genes Brain Behav, 10, 852–861.

McClung, C.A. (2007) Role for the Clock gene in bipolar disorder. Cold Spring Harb Symp Quant Biol, 72, 637–644.

McClung, C.A., Sidiropoulou, K., Vitaterna, M., Takahashi, J.S., White, F.J., Cooper, D.C. & Nestler, E.J. (2005) Regulation of dopaminergic transmission and cocaine reward by the Clock gene. Proc Natl Acad Sci U S A, 102, 9377–9381.

McGrath, C.L., Glatt, S.J., Sklar, P., Le-Niculescu, H., Kuczenski, R., Doyle, A.E., Biederman, J., Mick, E., Faraone, S.V., Niculescu, A.B. & Tsuang, M.T. (2009) Evidence for genetic association of RORB with bipolar disorder. BMC Psychiatry, 9, 70.

Meng, Q.J., Logunova, L., Maywood, E.S., Gallego, M., Lebiecki, J., Brown, T.M., Sladek, M., Semikhodskii, A.S., Glossop, N.R., Piggins, H.D., Chesham, J.E., Bechtold, D.A., Yoo, S.H., Takahashi, J.S., Virshup, D.M., Boot-Handford, R.P., Hastings, M.H. & Loudon, A.S. (2008) Setting clock speed in mammals: the CK1 epsilon tau mutation in mice accelerates circadian pacemakers by selectively destabilizing PERIOD proteins. Neuron, 58, 78–88.

Nesse, R.M. (1994) Fear and fitness: An evolutionary analysis of anxiety disorders. Ethology and sociobiology, 15, 247–261.

Nievergelt, C.M., Kripke, D.F., Barrett, T.B., Burg, E., Remick, R.A., Sadovnick, A.D., McElroy, S.L., Keck, P.E., Jr., Schork, N.J. & Kelsoe, J.R. (2006) Suggestive evidence for association of the circadian genes PERIOD3 and ARNTL with bipolar disorder. Am J Med Genet B Neuropsychiatr Genet, 141B, 234–241.

Oprea, T.I., Bologa, C.G., Brunak, S., Campbell, A., Gan, G.N., Gaulton, A., Gomez, S.M., Guha, R., Hersey, A., Holmes, J., Jadhav, A., Jensen, L.J., Johnson, G.L., Karlson, A., Leach, A.R., Ma’ayan, A., Malovannaya, A., Mani, S., Mathias, S.L., McManus, M.T., Meehan, T.F., von Mering, C., Muthas, D., Nguyen, D.T., Overington, J.P., Papadatos, G., Qin, J., Reich, C., Roth, B.L., Schürer, S.C., Simeonov, A., Sklar, L.A., Southall, N., Tomita, S., Tudose, I., Ursu, O., Vidovic, D., Waller, A., Westergaard, D., Yang, J.J. & Zahoránszky-Köhalmi, G. (2018) Unexplored therapeutic opportunities in the human genome. Nature reviews. Drug discovery, 17, 377.

Ouyang, Y., Andersson, C.R., Kondo, T., Golden, S.S. & Johnson, C.H. (1998) Resonating circadian clocks enhance fitness in cyanobacteria. Proceedings of the National Academy of Sciences, 95, 8660–8664.

Parker, G. & Brotchie, H. (2010) Gender differences in depression. International Review of Psychiatry, 22, 429–436.

Partonen, T., Treutlein, J., Alpman, A., Frank, J., Johansson, C., Depner, M., Aron, L., Rietschel, M., Wellek, S., Soronen, P., Paunio, T., Koch, A., Chen, P., Lathrop, M., Adolfsson, R., Persson, M.L., Kasper, S., Schalling, M., Peltonen, L. & Schumann, G. (2007) Three circadian clock genes Per2, Arntl, and Npas2 contribute to winter depression. Ann Med, 39, 229–238.

Patke, A., Young, M.W. & Axelrod, S. (2019) Molecular mechanisms and physiological importance of circadian rhythms. Nature Reviews Molecular Cell Biology, 1–18.

Pittendrigh, C., Bruce, V. & Withrow, R. (1959) Photoperiodism and related phenomena in plants and animals. Washington, DC: American Association for the Advancement of Science, 475–505.

Pittendrigh, C.S. & Minis, D.H. (1972) Circadian Systems: Longevity as a Function of Circadian Resonance in <em>Drosophila melanogaster</em>. Proceedings of the National Academy of Sciences, 69, 1537–1539.

Ponder, C.A., Kliethermes, C.L., Drew, M.R., Muller, J., Das, K., Risbrough, V.B., Crabbe, J.C., Gilliam, T.C. & Palmer, A.A. (2007a) Selection for contextual fear conditioning affects anxiety-like behaviors and gene expression. Genes Brain Behav, 6, 736–749.

Ponder, C.A., Munoz, M., Gilliam, T.C. & Palmer, A.A. (2007b) Genetic architecture of fear conditioning in chromosome substitution strains: relationship to measures of innate (unlearned) anxiety-like behavior. Mamm Genome, 18, 221–228.

Porcu, A., Vaughan, M., Nilsson, A., Arimoto, N., Lamia, K. & Welsh, D.K. (2020) Vulnerability to helpless behavior is regulated by the circadian clock component CRYPTOCHROME in the mouse nucleus accumbens. Proceedings of the National Academy of Sciences.

Prendergast, B.J., Onishi, K.G. & Zucker, I. (2014) Female mice liberated for inclusion in neuroscience and biomedical research. Neuroscience and biobehavioral reviews, 40, 1–5.

Prickaerts, J., Moechars, D., Cryns, K., Lenaerts, I., van Craenendonck, H., Goris, I., Daneels, G., Bouwknecht, J.A. & Steckler, T. (2006) Transgenic mice overexpressing glycogen synthase kinase 3beta: a putative model of hyperactivity and mania. J Neurosci, 26, 9022–9029.

Quiroz, J.A., Gould, T.D. & Manji, H.K. (2004) Molecular effects of lithium. Mol Interv, 4, 259–272.

R Development Core Team (2011) R: A language and environment for statistical computing. R Foundation for Statistical Computing, Vienna, Austria.

Ralph, M.R. & Menaker, M. (1988) A mutation of the circadian system in golden hamsters. Science (New York, N.Y.), 241, 1225–1227.

Richardson, M.P., Strange, B.A. & Dolan, R.J. (2004) Encoding of emotional memories depends on amygdala and hippocampus and their interactions. Nat Neurosci, 7, 278–285.

Roybal, K., Theobold, D., Graham, A., DiNieri, J.A., Russo, S.J., Krishnan, V., Chakravarty, S., Peevey, J., Oehrlein, N., Birnbaum, S., Vitaterna, M.H., Orsulak, P., Takahashi, J.S., Nestler, E.J., Carlezon, W.A., Jr. & McClung, C.A. (2007) Mania-like behavior induced by disruption of CLOCK. Proc Natl Acad Sci U S A, 104, 6406–6411.

Schwartz, G.E. & Weinberger, D.A. (1980) Patterns of emotional responses to affective situations: Relations among happiness, sadness, anger, fear, depression, and anxiety. Motivation and Emotion, 4, 175–191.

Severino, G., Manchia, M., Contu, P., Squassina, A., Lampus, S., Ardau, R., Chillotti, C. & Del Zompo, M. (2009) Association study in a Sardinian sample between bipolar disorder and the nuclear receptor REV-ERBalpha gene, a critical component of the circadian clock system. Bipolar Disord, 11, 215–220.

Shi, J., Wittke-Thompson, J.K., Badner, J.A., Hattori, E., Potash, J.B., Willour, V.L., McMahon, F.J., Gershon, E.S. & Liu, C. (2008) Clock genes may influence bipolar disorder susceptibility and dysfunctional circadian rhythm. Am J Med Genet B Neuropsychiatr Genet, 147B, 1047–1055.

Sokoloff, G., Parker, C.C., Lim, J.E. & Palmer, A.A. (2011) Anxiety and fear in a cross of C57BL/6J and DBA/2J mice: mapping overlapping and independent QTL for related traits. Genes, Brain and Behavior, 10, 604–614.

Soria, V., Martinez-Amoros, E., Escaramis, G., Valero, J., Perez-Egea, R., Garcia, C., Gutierrez-Zotes, A., Puigdemont, D., Bayes, M., Crespo, J.M., Martorell, L., Vilella, E., Labad, A., Vallejo, J., Perez, V., Menchon, J.M., Estivill, X., Gratacos, M. & Urretavizcaya, M. (2010) Differential association of circadian genes with mood disorders: CRY1 and NPAS2 are associated with unipolar major depression and CLOCK and VIP with bipolar disorder. Neuropsychopharmacology, 35, 1279–1289.

Spoelstra, K., Wikelski, M., Daan, S., Loudon, A.S. & Hau, M. (2016) Natural selection against a circadian clock gene mutation in mice. Proceedings of the National Academy of Sciences, 113, 686–691.

Stacklies, W., Redestig, H., Scholz, M., Walther, D. & Selbig, J. (2007) pcaMethods--a bioconductor package providing PCA methods for incomplete data. Bioinformatics, 23, 1164–1167.

Svenningsson, P., Tzavara, E.T., Carruthers, R., Rachleff, I., Wattler, S., Nehls, M., McKinzie, D.L., Fienberg, A.A., Nomikos, G.G. & Greengard, P. (2003) Diverse psychotomimetics act through a common signaling pathway. Science (New York, N. Y.), 302, 1412–1415.

Terracciano, A., Tanaka, T., Sutin, A.R., Sanna, S., Deiana, B., Lai, S., Uda, M., Schlessinger, D., Abecasis, G.R., Ferrucci, L. & Costa, P.T., Jr. (2010) Genomewide association scan of trait depression. Biological psychiatry, 68, 811–817.

Turek, F.W. (2007) From circadian rhythms to clock genes in depression. Int Clin Psychopharmacol, 22 Suppl 2, S1–8.

Vitaterna, M.H., King, D.P., Chang, A.M., Kornhauser, J.M., Lowrey, P.L., McDonald, J.D., Dove, W.F., Pinto, L.H., Turek, F.W. & Takahashi, J.S. (1994) Mutagenesis and mapping of a mouse gene, Clock, essential for circadian behavior. Science (New York, N.Y.), 264, 719–725.

Vitaterna, M.H., Ko, C.H., Chang, A.-M., Buhr, E.D., Fruechte, E.M., Schook, A., Antoch, M.P., Turek, F.W. & Takahashi, J.S. (2006) The mouse Clock mutation reduces circadian pacemaker amplitude and enhances efficacy of resetting stimuli and phase–response curve amplitude. Proceedings of the National Academy of Sciences, 103, 9327–9332.

von Saint Paul, U. & Aschoff, J. (1978) Longevity among blowfliesPhormia terraenovae RD kept in non-24-hour light-dark cycles. Journal of comparative physiology, 127, 191–195.

Woelfle, M.A., Ouyang, Y., Phanvijhitsiri, K. & Johnson, C.H. (2004) The adaptive value of circadian clocks: an experimental assessment in cyanobacteria. Current Biology, 14, 1481–1486.

Zhang, L., Hirano, A., Hsu, P.-K., Jones, C.R., Sakai, N., Okuro, M., McMahon, T., Yamazaki, M., Xu, Y., Saigoh, N., Saigoh, K., Lin, S.-T., Kaasik, K., Nishino, S., Ptácek, L.J. & Fu, Y.-H. (2016) A PERIOD3 variant causes a circadian phenotype and is associated with a seasonal mood trait. Proceedings of the National Academy of Sciences, 113, E1536–E1544.

Zheng, B., Albrecht, U., Kaasik, K., Sage, M., Lu, W., Vaishnav, S., Li, Q., Sun, Z.S., Eichele, G., Bradley, A. & Lee, C.C. (2001) Nonredundant roles of the mPer1 and mPer2 genes in the mammalian circadian clock. Cell, 105, 683–694.

Zhou, M., Rebholz, H., Brocia, C., Warner-Schmidt, J.L., Fienberg, A.A., Nairn, A.C., Greengard, P. & Flajolet, M. (2010) Forebrain overexpression of CK1delta leads to down-regulation of dopamine receptors and altered locomotor activity reminiscent of ADHD. Proc Natl Acad Sci U S A, 107, 4401–4406.

